# A systems chemoproteomic analysis of acyl-CoA/protein interaction networks

**DOI:** 10.1101/665281

**Authors:** Michaella J. Levy, David C. Montgomery, Mihaela E. Sardiu, Sarah E. Bergholtz, Kellie D. Nance, Jose Montano, Abigail L. Thorpe, Stephen D. Fox, Qishan Lin, Thorkell Andresson, Laurence Florens, Michael P. Washburn, Jordan L. Meier

**Author notes:** M.J.L. and D.C.M. contributed equally to this work.

## Abstract

Acyl-CoA/protein interactions are required for many functions essential to life including membrane synthesis, oxidative metabolism, and macromolecular acetylation. However, despite their importance, the global scope and selectivity of these protein-metabolite interactions remains undefined. Here we describe the development of CATNIP (CoA/AcetylTraNsferase Interaction Profiling), a chemoproteomic platform for the high-throughput analysis of acyl-CoA/protein interactions in endogenous proteomes. First, we apply CATNIP to identify acetyl-CoA-binding proteins through unbiased clustering of competitive dose-response data. Next, we use this method to profile diverse protein-CoA metabolite interactions, identifying biological processes susceptible to altered acetyl-CoA levels. Finally, we apply systems-level analyses to assess the features of novel protein networks that may interact with acyl-CoAs, and demonstrate a strategy for high-confidence proteomic annotation of acetyl-CoA binding proteins. Overall our studies illustrate the power of integrating chemoproteomics and systems biology, and provide a resource for understanding the roles of acyl-CoA metabolites in biology and disease.

## Introduction

Acyl-CoAs are essential for life. These metabolites serve as fundamental cellular building blocks in the biosynthesis of lipids, intermediates in energy production via the TCA cycle, and essential precursors for reversible protein acetylation. Each of these functions are physically dependent on acyl-CoA/protein interactions, which can regulate protein activity via a variety of mechanisms (Fig. 1). For example, the interaction of acyl-CoAs with lysine acetyltransferase (KAT) active sites allows them to serve as enzyme cofactors or, alternatively, competitive inhibitors (Dyda et al., 2000; Montgomery et al., 2015). Binding of acyl-CoAs to the allosteric site of pantothenate kinase (PanK) enzymes can exert positive or negative effects on CoA biosynthesis (Hong et al., 2007). Acyl-CoAs can also non-enzymatically modify proteins, a covalent interaction that often causes enzyme inhibition (Kulkarni et al., 2017; Wagner et al., 2017). These examples illustrate the ability of acyl-CoA signaling to influence biology and disease. However, the global scope and selectivity of these metabolite-governed regulatory networks remains unknown.

**Figure 1.**
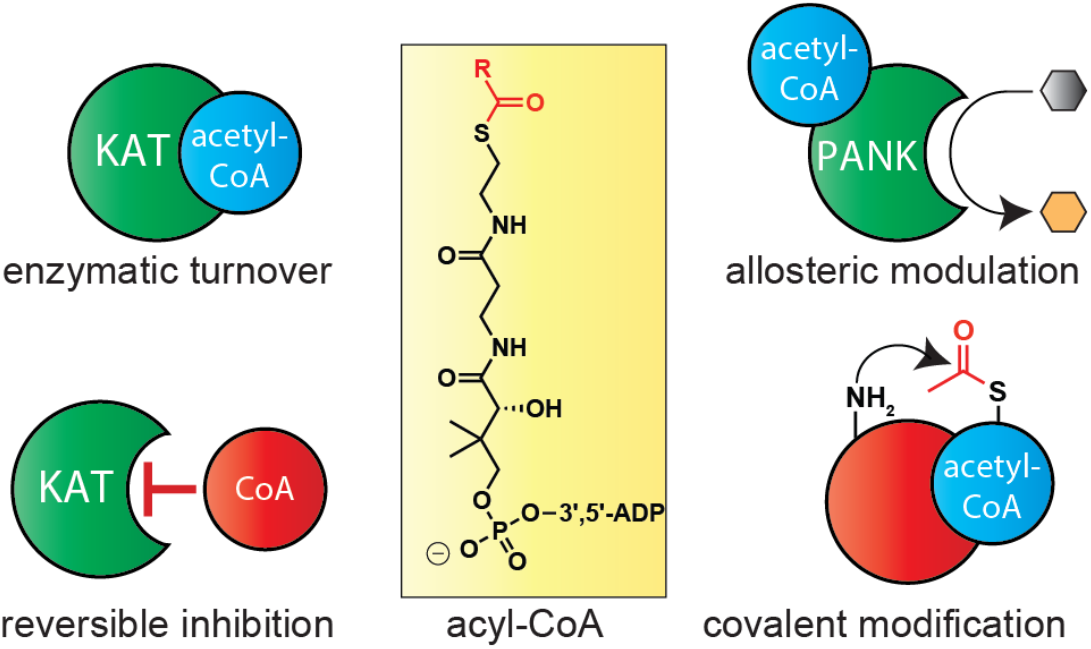
Diverse consequences of acyl-CoA interactions on protein activity and signaling. Metabolic acyl-CoAs can interact with proteins as enzymatic cofactors (top left), reversible inhibitors (bottom left), allosteric modulators (top right), or covalent modifiers (bottom right).

A central challenge of studying acyl-CoA/protein interactions is their pharmacological nature (Kulkarni et al., 2019). These transient binding events are invisible to traditional next-generation sequencing and proteomic methods. To address this, our group recently reported a competitive chemical proteomic (“chemoproteomic”) approach to detect and analyze acyl-CoA/protein binding (Montgomery et al., 2016; Montgomery et al., 2014). This method applies a resin-immobilized CoA analogue (Lys-CoA) as an affinity matrix to capture CoA-utilizing enzymes directly from biological samples. Pre-incubating proteomes with acyl-CoA metabolites competes capture and allows their relative binding affinities to enzymes of interest to be assessed. In our initial application of this platform we studied the susceptibility of KATs to metabolic feedback inhibition by CoA, evaluating competition by quantitative immunoblot (Montgomery et al., 2016). The signal amplification afforded by immunodetection enables the capture of extremely low abundance KATs to be readily quantified; however, due to its targeted nature, it is best suited to the study of specific CoA-utilizing enzymes rather than broad profiling or discovery applications. We reasoned such applications could be enabled by integrating CoA-based affinity reagents with i) multidimensional chromatographic separation, to efficiently sample rare KAT enzymes, ii) quantitative LC-MS/MS proteomics, for unbiased identification of CoA-interacting proteins, and iii) systems analysis of the acyl-CoA-binding proteins identified, for data-driven analysis of putative interaction networks. We term this approach CATNIP (<u>C</u>oA/<u>A</u>cetyl<u>T</u>ra<u>N</u>sferase <u>I</u>nteraction <u>P</u>rofiling). Here we describe the development and application of CATNIP to globally analyze acyl-CoA/protein interactions in endogenous human proteomes. First, we demonstrate the ability of CATNIP to identify acetyl-CoA-binding proteins through unbiased clustering of competitive dose-response data. Next, we apply this method to profile diverse protein-CoA metabolite interactions, enabling the identification of biological processes susceptible to altered acetyl-CoA levels. Finally, we utilize systems-level analyses to assess the features of novel protein networks that may interact with acyl-CoAs and demonstrate a strategy for high-confidence annotation of direct acetyl-CoA binding proteins and AT enzymes in human proteomes. Overall our studies illustrate the power of integrating chemoproteomics and systems biology analysis methods and provide a novel resource for understanding the diverse signaling roles of acyl-CoAs in biology and disease.

## Results

### Validation of CATNIP for the global study of acyl-CoA/protein interactions

In order to deeply sample acyl-CoA/protein interactions on a proteome-wide scale, we initially set out to integrate CoA-based protein capture methods with LC-MS/MS (Fig. 2a). In this workflow, whole cell extracts are first incubated with Lys-CoA Sepharose. This affinity matrix enables active site-dependent enrichment of many different classes of CoA-binding proteins (Montgomery et al., 2016), making it ideal for broad profiling studies. Next, enriched proteins are subjected to tryptic digest and analyzed using MudPIT (multidimensional protein identification technology), a proteomics platform that combines strong cation exchange and C18 reverse phase chromatography to pre-fractionate tryptic peptides, followed by ionization and data-dependent MS/MS (Washburn et al., 2001b). The separation afforded by this approach significantly decreases sample complexity, allowing the identification of rare, low abundance peptides from complex proteomic mixtures. To facilitate the identification of acyl-CoA/protein interactions, competition experiments are performed in which proteomes are pre-incubated with a CoA metabolite prior to capture (Leung et al., 2003). Decreased enrichment in competition samples compared to controls (as assessed by quantitative spectral counting) signifies that the CoA metabolite interacts with a protein of interest. These interacting proteins can then be further classified into pharmacological or biological networks using either conventional metrics (fold-change, gene ontology, etc) or systems-based analysis tools.

**Figure 2.**
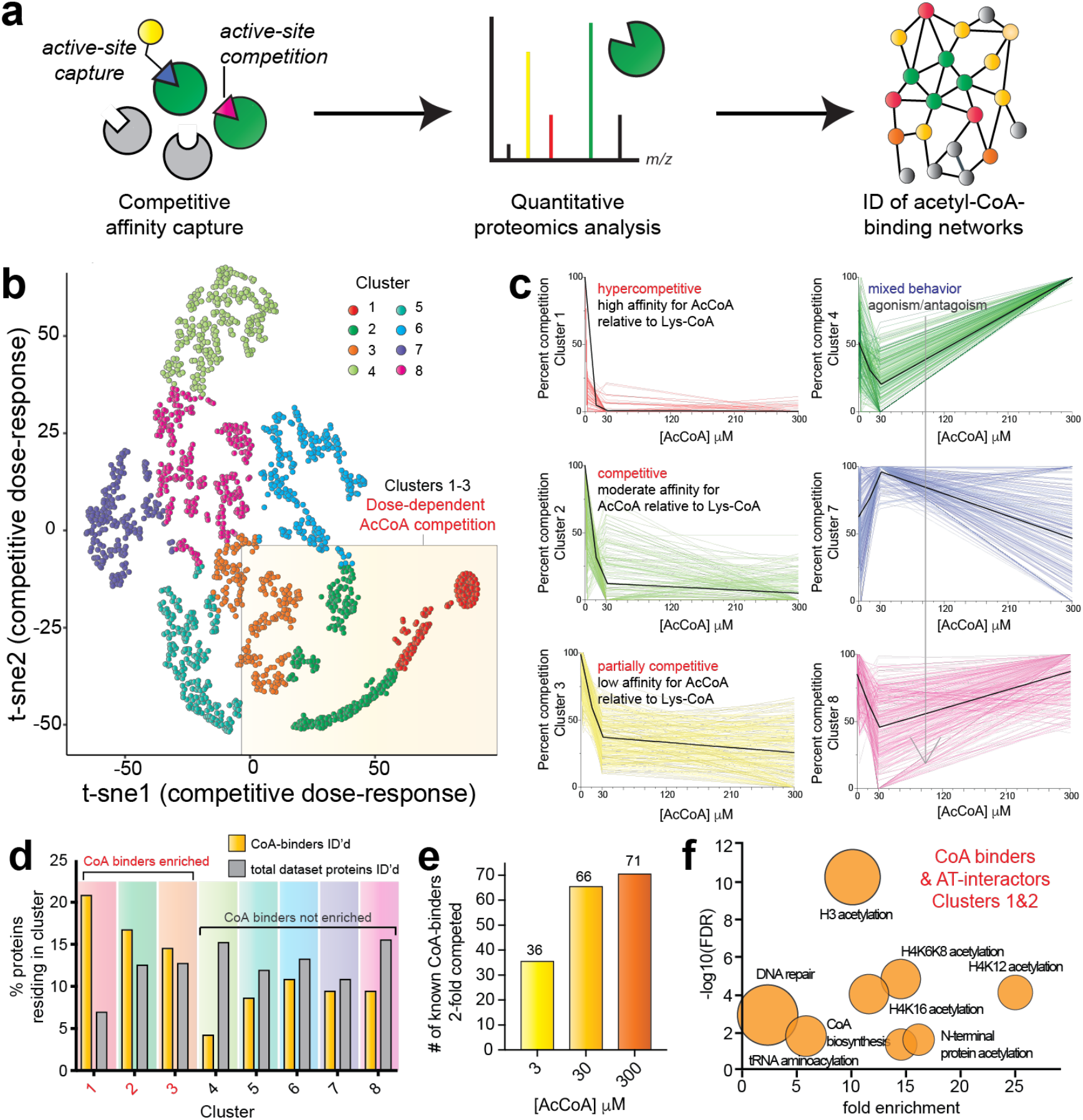
Profiling acetyl-CoA/protein interactions using CATNIP. (a) Schematic for chemoproteomic analyses of acyl-CoA/protein interactions. (b) Normalized dNSAF values across the 4 acetyl-CoA concentrations (0, 3, 30, 300μM) were t-SNE transformed and plotted in two dimensions for all proteins competed by the various concentrations of acetyl-CoA in the CATNIP experiments. Eight clusters were identified as the optimal number by k-means analysis. For all LC-MS/MS experiments, n=3. Each cluster is colored according to the legend.(c) Dose-response profiles of acetyl-CoA in the CATNIP clusters. Colored lines indicate the capture profiles of individual proteins within each cluster in the presence of increasing concentrations of acetyl-CoA competitor. Black lines indicate the mean capture profile for all proteins in a given cluster. (d) Acetyl-CoA competitive clusters 1-3 are enriched in Uniprot-annotated CoA-binding proteins (“CoA binders”) as well as members of acetyltransferase complexes (”AT interactors”). (e) The number of annotated CoA binders exhibiting 2-fold competition in the presence 30 and 300 HM competition is similar. (f) Gene ontology analysis of Uniprot annotated CoA-binding and AT interacting proteins lying in CATNIP clusters 1 and 2. Fold enrichment of a specific functional term is plotted versus statistical significance (-log10[FDR]). The circle size reflects the number of proteins matching a given term. Functional enrichment was performed with the tool DAVID (https://david.ncifcrf.gov) by using GO and Swiss-Prot Protein Information Resource terms.

As an initial model, we explored the utility of CATNIP to globally profile acetyl-CoA/protein interactions in unfractionated HeLa cell proteomes. Proteomes were pre-incubated with acetyl-CoA or vehicle (buffer) control, followed by enrichment using Lys-CoA Sepharose. These experiments assessed competition at 3, 30, and 300 μM acetyl-CoA, which spans the physiological concentration range of acetyl-CoA in the cytosol and mitochondria. Protein capture in each condition was quantified using distributed normalized spectral abundance factor (dNSAF), a label-free metric that normalizes spectral counts relative to overall protein length (Fig. S1a-c, Table S1) (Zhang et al., 2015). Each condition was analyzed in triplicate, constituting 12 experiments, >144 hours of instrument time, and over 1.1 million non-redundant peptide spectra collected. We limited our analysis to high-confidence protein identifications (>4 spectral counts in vehicle [0 μM] sample). The capture of Uniprot annotated CoA-binding proteins or members of AT complexes (termed ‘AT interactors’) did not correlate with overall protein abundance or gene expression (Nagaraj et al., 2011), consistent with the ability of chemoproteomic methods to sample functional activity metrics (e.g. unique pharmacology, active-site folding/conformation, integration into complexes, posttranslational modification) rather than raw quantity (Fig. S1d-i) (Moellering and Cravatt, 2012).

To analyze acetyl-CoA binding in a systematic manner throughout the proteome, we first grouped proteins into subsets based on their dose-dependent competition profiles. Chemoproteomic capture data from 0, 3, 30, and 300 μM acetyl-CoA competition was transformed, plotted in two dimensions, and subjected to k-means clustering. Eight protein clusters were identified, each of which exhibited a distinct dose-dependent competition signature (Fig. 2b-c, Fig. S2a-b). The capture of proteins within clusters 1-3 were antagonized by acetyl-CoA in a dose-dependent fashion. Proteins in cluster 1 displayed hypersensitivity to acetyl-CoA competition, while proteins in clusters 2 and 3 exhibited moderate and partial competition, respectively. The remaining clusters exhibited more complicated capture profiles, consisting of either dose-dependent and independent antagonism (cluster 5), or mixed agonist/antagonist behavior (clusters 4, 6-8, Fig. 2c, Fig. S2b). To determine which of these competition signatures were most characteristic of acetyl-CoA binding, we first analyzed each cluster for the presence of known CoA-binding proteins and AT interactors. Cluster 1, composed of proteins whose capture is hypercompetitive to pre-incubation with acetyl-CoA, contains only 7% of the total proteins identified in this experiment. However, 25% of proteins in this cluster are annotated CoA-binding proteins and AT interactors, a disproportionate enrichment (Fig. 2d, Fig. S2c). Clusters 2 and 3 were also relatively enriched in annotated CoA binders and AT interactors, while all other subsets were not (Fig. 2d). Examining our entire dataset, we found the total number of CoA-binding proteins and AT interactors competed 2-fold by acetyl-CoA almost doubled going from 3 to 30 μM, but was only modestly increased by higher concentrations of competitor (Fig. 2e). Proteins in clusters 1 and 2 exhibit almost complete loss of capture at 30 μM acetyl-CoA (Fig. 2c). This suggests the occupancy of most acetyl CoA-binding sites accessible to our method are saturated at the intermediate concentration used here (~30 μM), in line with literature measurements of binding affinity and Michaelis constants (Scheer et al., 2011). Clusters 1 and 2 include proteins that bind to acetyl-CoA directly (CREBBP, NAA10), allosterically (PANK1), and indirectly via protein-protein interactions (NAA25, JADE1; Table S2). This indicates that proteins with disparate modes of acetyl-CoA interaction can display similar dose-dependent competition signatures. Gene ontology analysis of annotated CoA binders in clusters 1 and 2, whose enrichment was hypercompetitive to acetyl-CoA pre-incubation, revealed an enrichment in terms related to histone and N-terminal acetyltransferases as well as CoA biosynthetic enzymes (Fig. 2f, Fig. S2d-e). The strong enrichment of KATs likely results from the propensity of our bisubstrate Lys-CoA capture agent to interact with this enzyme family (Lau et al., 2000). A similar analysis of proteins in cluster 3, which exhibits partial competition by acetyl-CoA, identified a disproportionate number of mitochondrial CoA-binding enzymes (Fig. S2f). This decreased sensitivity to acyl-CoA competition may reflect evolutionary adaptation to the unique subcellular concentrations of metabolites in mitochondria, where acetyl-CoA is found at millimolar concentrations (Chen et al., 2016). Overall, these studies validate the ability of CATNIP to detect bona fide acetyl-CoA/protein interaction signatures, and establish key parameters necessary for its useful application in studying the pharmacology of CoA metabolites.

### Applying CATNIP to define the unique pharmacological signatures of acetyltransferase enzymes

In addition to acetyl-CoA (**1**), cells produce a physiochemically diverse range of CoA metabolites whose concentrations directly reflect the metabolic state of the cell. Many of these species make regulatory interactions with proteins, including the long chain fatty acyl (LCFA) palmitoyl-CoA, a classic feedback inhibitor of acetyl-CoA carboxylase (Greenspan and Lowenstein, 1968), short chain fatty acyl (SCFA) butyryl-CoA, which can potently inhibit KATs or be used as a substrate (Carrer et al., 2017; Montgomery et al., 2015), and negatively charged succinyl-CoA, which can covalently inhibit many mitochondrial enzymes (Kulkarni et al., 2017; Wagner et al., 2017). However, despite their physiological relevance, few studies have interrogated the comparative pharmacology of acyl-CoA/enzyme interactions. We hypothesized that the ability of CATNIP to report on the binding affinity of ligands relative to Lys-CoA could address this gap and enable the generation of pharmacological fingerprints of acyl-CoA-protein interactions across the proteome. To explore this hypothesis, we performed competitive chemoproteomic capture experiments in the presence of additional metabolites including: i) CoA (**2**), a feedback inhibitor of acetyltransferases, ii) butyryl-CoA (**3**), a short chain fatty acyl-CoA, iii) crotonyl-CoA (**4**), a SCFA-CoA containing a latent acrylamide electrophile, iv) acetic-CoA (**5**), a stable analogue of malonyl-CoA which has recently been shown to be a hyperreactive metabolite capable of covalent protein modification, and v) palmitoyl-CoA (**6**), a LCFA-CoA which we have previously shown can potently inhibit KATs in vitro (Fig. 3a, Table S3). For these experiments, CoA metabolites were equilibrated with proteomes (1 h) prior to Lys-CoA capture. A dosage of 30 μM was selected to enable a comparison of each ligand’s competition profile to that of acetyl-CoA, which showed substantial interaction with proteins in clusters 1-3 at this concentration. For palmitoyl-CoA a lower concentration was used (3 μM) in order to ensure solubility and reflect the limited free (non-protein/membrane bound) quantities of LCFA-CoA likely to be present in cells.

**Figure 3.**
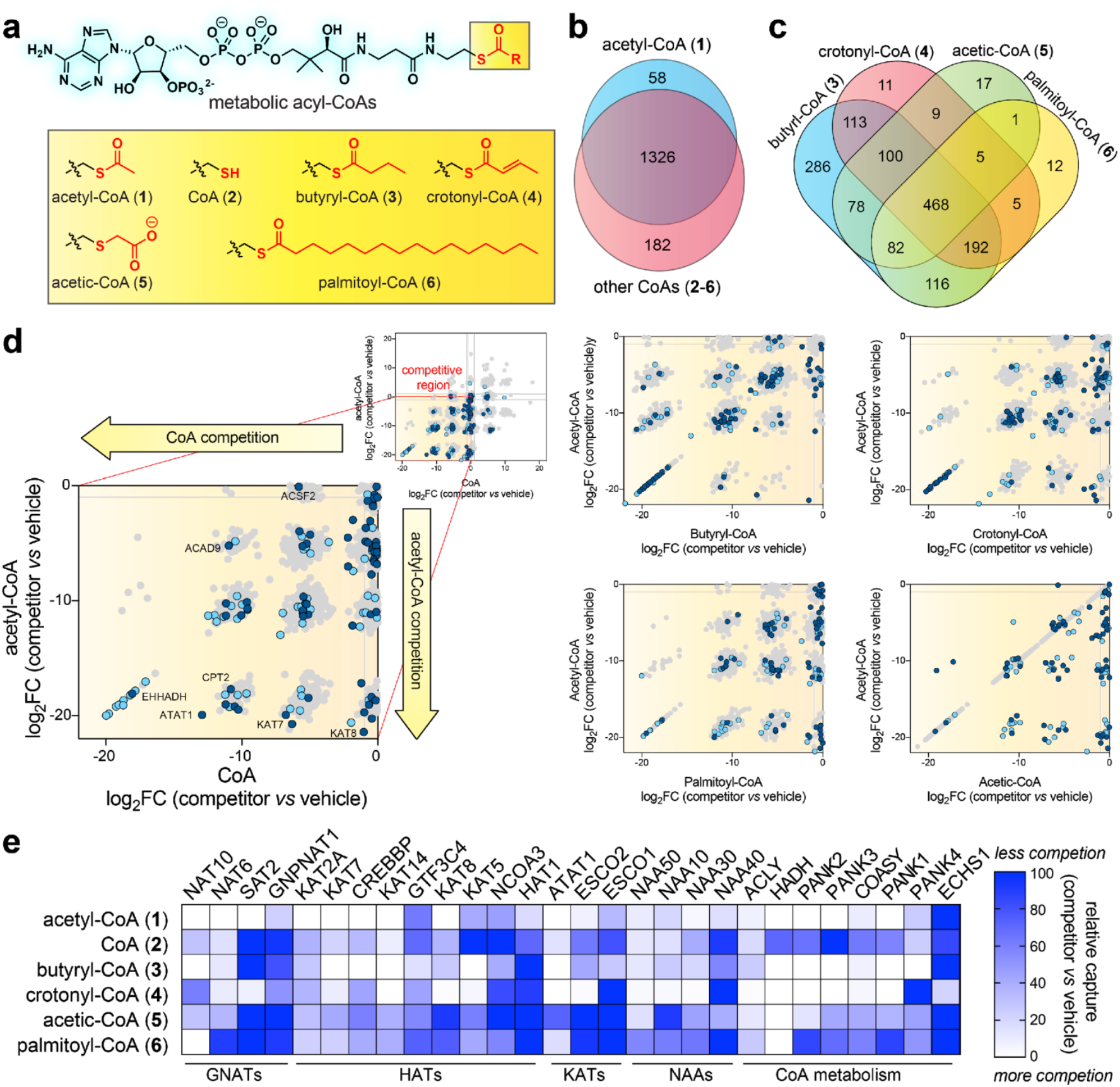
Applying CATNIP to profile the comparative pharmacology of diverse CoA metabolites. (a) CoA metabolites (**1**-**6**) analyzed in this study. (b) Venn diagram depicting overlap between proteins whose capture by Lys-CoA Sepharose was competed more than two-fold by acetyl-CoA (**1**) or all other CoAs (**2-6**). (c) Venn diagram depicting overlap between proteins whose capture by competed by acyl-CoAs **3**-**6**. (d) Comparison of acetyl-CoA and acyl-CoA/protein interaction profiles. Proteins are plotted according to competition of capture by each metabolic acyl-CoA (x-axis) and acetyl-CoA (y-axis). Uniprot-annotated CoA-binding proteins (“CoA binders”) as well as members of acetyltransferase complexes (”AT interactors”) are highlighted in dark blue and light blue, respectively. Competition data for additional doses of CoA (**2**) can be found in the Supporting Information. (e) Comparative CATNIP analysis indicates proteins from related protein families can display distinctive signatures of pharmacological interaction with CoA metabolites. White = more competition by metabolic acyl-CoA, blue = less competition by metabolic acyl-CoA.

As an initial rough measure of acyl-CoA selectivity, we performed a global analysis of proteins displaying robust interaction (>2-fold decreased capture) with competitors. Evaluating 1757 proteins quantified in Lys-CoA enrichments, we found that 1566 (89%) were >2-fold competed by at least one CoA/acyl-CoA metabolite (**1**-**6**, Fig. 3b, Table S3). In general the majority of proteins found to interact with acetyl-CoA (**1**) also displayed competition by **2**-**6**, suggestive of ligand-binding promiscuity amongst CoA-binding proteins. Examining physiochemically distinct ligands **3**-**6**, a handful of selective interactions were observed for each acyl-CoA (Fig. 3c). Notably, butyryl-CoA (**3**) showed substantial overlap with protein interactors of **4**-**6**. This may be suggestive of its metabolic stability in lysates, or ability to make high affinity interactions with many classes of CoA-binding proteins at the concentration applied. To compare the magnitude of protein-ligand interactions, we plotted the competition (log2 fold change, competitor v. control) of each individual ligand relative to acetyl-CoA (Fig. 3d). Most proteins interacted more strongly with acetyl-CoA (**1**) than other ligands, with the exception of butyryl-CoA (**3**). This is consistent with the fact that acetyl-CoA (**1**) and butyryl-CoA (**3**) exhibited the greatest number of unique interaction partners in our comparative analysis. This promiscuous binding also extended to known CoA-binding proteins, including KATs (Fig. S3). Notable exceptions were HADHB, which was found to interact only with butyryl-CoA, as well as ECHS1, which was found to interact only with crotonyl-CoA (Table S3). HADHB encodes the thiolase subunit of the mitochondrial trifunctional protein, which is involved in the oxidation of fatty acids 8 carbons or longer (Middleton, 1994). The ability of this enzyme to specifically interact with butyryl-CoA but not acetyl-CoA could represent a mechanism allowing cells to sense blockade of the terminal steps of SCFA-CoA catabolism, triggering feedback inhibition of fatty acid oxidation in a manner that product inhibition does not. ECHS1 encodes an acyl-CoA dehydrogenase, and was the only protein found to be competed 2-fold by crotonyl-CoA, but not the structurally related acetyl- and butyryl-CoA. Other enzymes with crotonase folds did not display this selective inhibition profile (Table S4). Selective crotonyl-CoA dependent capture is consistent with the substrate specificity of this enzyme, which shows rapid turnover of crotonyl-CoA relative to longer chain enoyl-CoA thioesters (Yamada et al., 2015). To facilitate a more granular analysis, we grouped CoA-binding proteins by biological function or fold and compared their quantitative metabolite-binding signatures upon interaction with **1**-**6**. Histone, lysine, and GNAT acetyltransferases displayed a diversity of ligand binding signatures (Fig. 3e). For example, the capture of enzymes such as CREBBP and NAT10 was strongly competed by multiple metabolites, while others (KAT8 and HAT1) displayed an apparent preference for selective interaction with acetyl-CoA (Fig. 3e). Selectivity did not correlate with enzyme function/fold, capture abundance, or acetyl-CoA interaction cluster (Fig. 3e, Table S4), suggesting this metabolite interaction fingerprint represents a unique and intrinsic feature of individual enzymes. The promiscuous ligand binding of CREBBP is notable, as this KAT and its homologue EP300 have been found to utilize several acyl-CoAs as alternative cofactors (Chen et al., 2007; Sabari et al., 2018). The PANK family of proteins catalyze the phosphorylation of pantothentate (vitamin B5) to phosphopantothenate, which constitutes a key step in CoA biosynthesis. Previous biochemical studies have found PANK1 to be allosterically inhibited by acetyl-CoA but not CoA, while PANK2 is strongly inhibited by both ligands (Rock et al., 2002; Zhang et al., 2006). We found acetyl-CoA interacted more strongly with each enzyme, but did not observe substantial disparity between CoA binding to the two enzyme isoforms. This may reflect differential binding of metabolites to these enzymes in the complex proteomic milieu compared to biochemical assays or, alternatively, a limitation of our method, which uses a single concentration of ligand that may saturate both selective and non-selective interactions. Overall, these studies validate the ability of chemoproteomics to study acyl-CoA/protein interactions and

### Evaluating the dynamic activity of acetyltransferases in response to metabolic perturbation

Coenzyme A (**2**) is one of the most abundant metabolites in cells. In addition to functioning as an obligate precursor for acyl-CoA biosynthesis, CoA can also serve as a potent feedback inhibitor of members of the AT superfamily. Previously, we used quantitative immunoblotting of chemoproteomic capture experiments to probe the sensitivity of eight ATs to product inhibition by testing their relative binding to acetyl-CoA (cofactor) and CoA (inhibitor) (Montgomery et al., 2016). The success of this approach inspired us to apply CATNIP to extend this comparison proteome-wide. Capture experiments were performed in the presence of escalating doses of CoA (3, 30, 300 μM), transformed, and clustered using an identical pipeline as in our acetyl-CoA binding experiments above (Table S5). Two clusters (3 and 8) exhibited readily interpretable dose-dependent competition profiles, with several additional clusters (1, 2, and 5) displaying hypersensitivity at low concentrations (3 μM) of CoA (Fig. S4a-b). Dose-dependent cluster 3 contained KAT2A, CREBBP, and PANK2, all of whom have been shown to be biologically or biochemically susceptible to metabolic feedback inhibition by CoA (Hong et al., 2007; Marino et al., 2014; Tanner et al., 2000). Further examination of this cluster revealed three proteins that were most sensitive to CoA, exhibiting >50% loss of capture in the presence of 3 μM ligand and >80% loss of capture in the presence of 30 μM ligand: ACLY, NAT6, and NAT10 (Fig. 4a). The unusually strong CoA interaction profile of these three proteins was distinct from that of other proteins in the cluster and within the KAT superfamily, most of which are competed much more efficiently by acetyl-CoA (Fig. 4b, Fig. S4c). The binding of ACLY to both CoA and acetyl-CoA is consistent with the reversible activity of the enzyme, which has been previously observed in biochemical assays (Inoue et al., 1968). NAT6 (NAA80) is a recently de-orphanized enzyme which has been determined to acetylate the N-terminus of actin, whose metabolic sensitivity has not been explored (Drazic et al., 2018). NAT10 is an RNA acetyltransferase that has been found to catalyze acetylation of cytidine in ribosomal, transfer, and messenger RNA, forming the minor nucleobase N4-acetylcytidine (ac4C) (Arango et al., 2018; Ito et al., 2014; Sharma et al., 2015). The identification of NAT10-CoA interactions by CATNIP is consistent with our previous studies, which have established that NAT10 binds acetyl-CoA and CoA with similar affinities and may be susceptible to metabolic feedback inhibition (Montgomery et al., 2016). Compounding this effect, no related N4-acylations of cytidine (e.g. butyrylation) have been identified in RNA, suggesting NAT10 may additionally interact with SCFA- and LCFA-CoAs as inhibitors.

**Figure 4.**
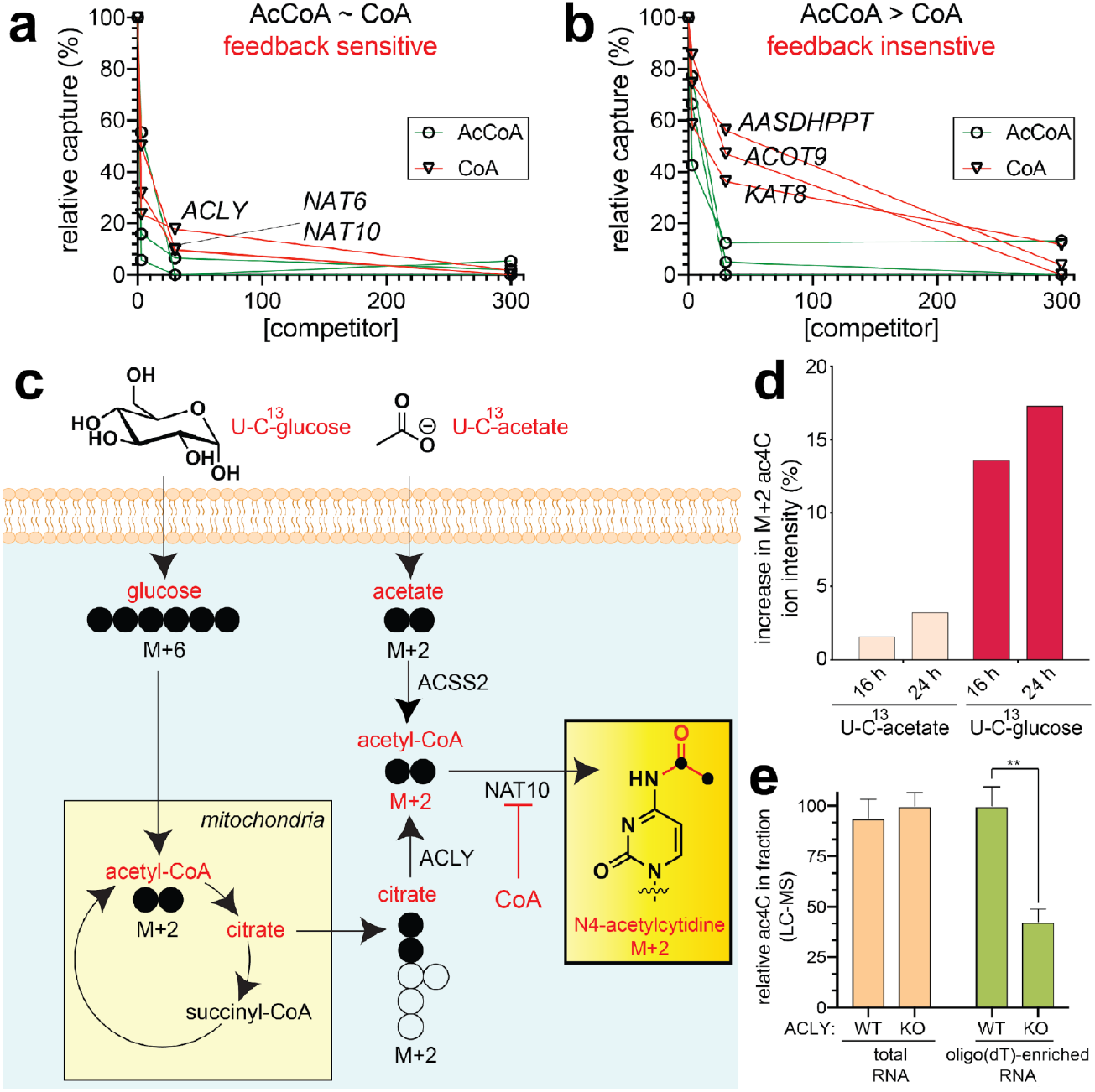
Applying CATNIP to profile the susceptibility of ATs to metabolic feedback inhibition. (a) Exemplary competitive dose-response profiles of proteins that interact strongly with CoA and acetyl-CoA. (b) Exemplary competitive dose-response profiles of proteins that interact moderately with CoA and strongly with acetyl-CoA. (c) Scheme for isotopic tracing experiments designed to determine the metabolic source of the acetate group in ac4C. Heavy (U-^13^C) glucose or acetate were applied in separate metabolic labeling experiments. Incorporation into ac4C was assessed by digest of total RNA to constituent nucleotides followed by mass isotopomer analysis. (d) Metabolic tracing reveals the major source of ac4C’s N4-acetyl group is glucose-derived acetyl-CoA. (e) Disruption of ACLY-dependent glucose-derived acetyl-CoA production reduces levels of ac4C in poly(A)-enriched, but not total RNA, fraction. Values represent ≥ 3 replicates, analyzed by two-tailed student’s t-test (ns = not significant, *P<0.05, **P<0.01, ***P<0.001).

To explore the metabolic inhibition of NAT10 in greater detail, we determined the metabolic source of the acetate group post-transcriptionally introduced into ac4C in proliferating cancer cell lines (Fig. 4c). Treatment of cells with isotopically-labeled acetyl-CoA precursors, followed by RNA digest and analysis by LC-MS/MS, revealed that the majority of NAT10-dependent cytidine acetylation stems from glucose-derived acetyl-CoA (Fig. 4d). Since the production of glucose-derived acetyl-CoA in human cells is highly dependent on ACLY activity, and ACLY perturbation can drastically influence the ratio of acetyl-CoA to inhibitory CoA metabolites,(Wellen et al., 2009) we next examined how stable knockout of ACLY impacted ac4C levels in RNA. Analysis of wild-type and ACLY knockout human glioblastoma cells (Zhao et al., 2016) revealed similar levels of ac4C in total RNA (Fig. 4e). However, LC-MS analysis of poly(A)-enriched RNA fractions from these cell lines indicated an ACLY-dependent decrease in ac4C. ACLY-dependent deposition is also observed for another acetyl-CoA derived RNA nucleobase, 5-methoxycarbonylmethyl-2-thiouridine (mcm5S2U), whose production is catalyzed by the AT enzyme Elp3 (Fig. S4d-e) (Lin et al., 2019). The observation that ac4C and mcm5s2U are sensitive to the metabolic state of the cell is consistent with the findings of Balasubramanian and coworkers, who reported that starvation conditions reduced NAT10-dependent ac4C levels in transfer RNA (van Delft et al., 2017). The ability of ACLY perturbation to influence the acetylation of poly(A)RNA, but not total RNA, suggests inhibitory CoA/acyl-CoAs may interact in a distinct manner with different functional forms of NAT10. These studies illustrate the ability of CATNIP to guide the identification of novel acetylation events that are sensitive to the metabolic state of the cell.

### Unbiased CATNIP analysis reveals annotation and mechanistic features of acyl-CoA-binding

The annotation of the cellular acyl-CoA binding proteome has never been directly assessed using experimental methods. Therefore, we next set out to develop an unbiased workflow for analysis of CATNIP binding data that could enable the de novo identification of known acyl-CoA dependent enzymes and ask what, if any, uncharacterized proteins share these properties. To differentiate acyl-CoA interacting proteins from background, our initial criteria were: 1) significant competition (*p* ≤ 0.05) of enriched proteins by three or more CoA ligands, and 2) absence of enriched proteins in the ‘CRAPome’ common contaminant database (Fig. 5a) (Mellacheruvu et al., 2013). Of 1764 proteins detected in Lys-CoA Sepharose capture experiments, 672 (38.1%) passed these cut-offs (Table S6), including the majority of annotated ATs that were enriched by Lys-CoA (Fig. 5b). Acetyltransferases not identified were mostly found to be poorly expressed by RNA Seq (Fig. 5b) (Nagaraj et al., 2011) and did not display obvious structural similarities in GNAT consensus elements (Fig. S5a) (Dyda et al., 2000). To examine whether unique patterns of acyl-CoA binding in this filtered dataset are associated with distinct biological processes, we further analyzed these proteins using Topological Data Analysis (TDA) (Lum et al., 2013). TDA functions as a geometric approach that can be used to identify shared properties of complex multidimensional datasets that may not be apparent by other methods, and has previously been used to detect biologically-relevant modules in protein complexes from immunoprecipitation LC-MS/MS data (Sardiu et al., 2015). Therefore, we applied TDA to analyze the multidimensional CoA metabolite competition profiles for each protein in our filtered subset, and then annotated the TDA clusters with enriched pathways identified by gene ontology analysis using DAVID (https://david.ncifcrf.gov) and ConsensusPathDB (http://cpdb.molgen.mpg.de/). This analysis revealed that histone acetyltransferases, which bind to CoAs directly, form a distinct cluster relative to PANK2 and PANK3, which are allosterically regulated by CoA metabolites. This analysis also identified many proteins involved in RNA metabolism and cell cycle whose association with CoA metabolites has not been previously characterized (Fig. 5c). This suggests that multiple proteins involved in these processes may directly or indirectly bind to acyl-CoAs, and potentially be subject to differential regulation CoA metabolism. Moreover, these studies demonstrate the utility of TDA for clustering and visual representation of ligand-protein interaction networks.

**Figure 5.**
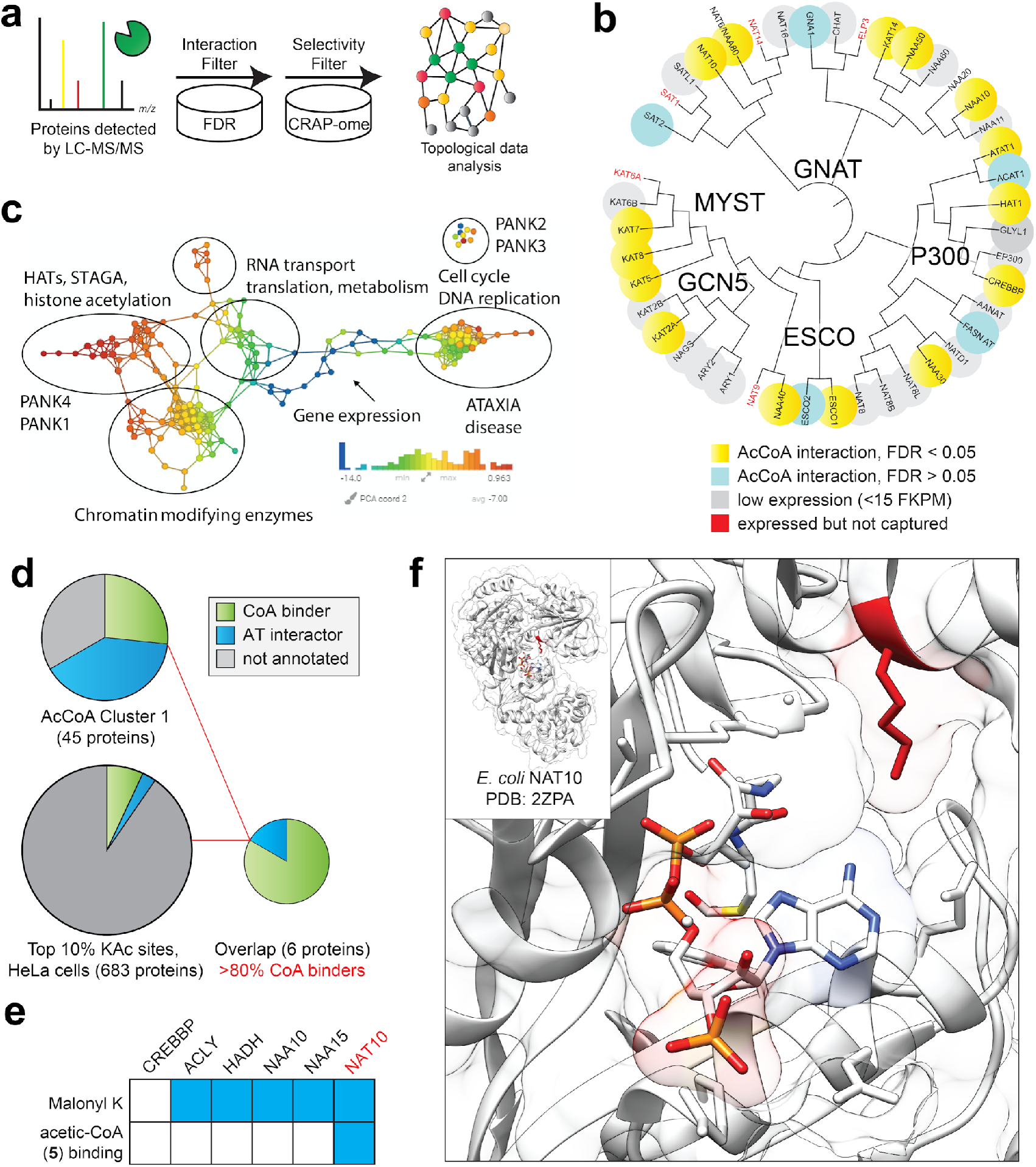
De novo annotation of the CoA-binding proteome. (a) Stringent filtering of CATNIP-enriched proteins using statistically significant competition by multiple (>3) ligands as well as absence from common contaminant databases was used as an initial metric to differentiate potential CoA-binders from background proteins. (b) Proteins from diverse protein, RNA, and metabolite AT families display statistically significant multi-ligand competition. The majority of proteins not detected were found to be poorly expressed in HeLa by RNA-Seq. (c) Topological network analysis of proteins exhibiting significant interaction with >3 CoA metabolites reveals overlapping networks of acyl-CoA protein-interactions. Protein nodes are colored based on the metric PCA2. Color bar: red = high values; blue = low values. Node size is proportional to the number of proteins in the node. (d) Combining multi-ligand CATNIP competition and acetylation stoichiometry filters greatly enriches CoA-binders and AT-interactors relative to either measure alone. (e) Multiple high-confidence CoA-binders detected by de novo CATNIP analysis contain annotated sites of lysine malonylation (top row). However, only NAT10 displays statistically significant competition of capture by malonyl-CoA mimic **5** (bottom row). (f) A conserved site of lysine malonylation lies in close proximity to the acetyl-CoA binding site of a bacterial NAT10 orthologue.

Next, we sought to examine the annotation of the human CoA-binding proteome. Specifically, we wished to incorporate additional criteria allowing us to differentiate proteins that directly bind to CoA metabolites, such as ATs, from proteins that are indirectly captured via protein-protein interactions, such as non-catalytic members of AT complexes. Such an approach would potentially provide a pipeline for novel AT discovery, as well as insights into how well the CoA-binding proteome is currently characterized. To accomplish this, we first classified proteins in our statistically significant filtered subset based on their dose-dependent acetyl-CoA competition profiles determined above, whose clustering we found could highlight protein subsets enriched in known ATs and CoA-binding proteins (Fig. 2b, d). Approximately 7% (45/672) of the filtered proteins resided in cluster 1, whose capture is hypersensitive to competition by acetyl-CoA (Fig. 5d, Table S7). This included 12 direct CoA-binders, 15 AT-interactors, and 18 proteins whose interaction with CoA metabolites had not been previously characterized. Amongst this protein subset, terms related to histone acetyltransferases and CoA biosynthesis were clearly differentiated as the most highly enriched biological process (Fig. S5d). To further differentiate direct and indirect CoA interactions, we assessed these 45 proteins for sites of high stoichiometry acetylation. Our reasoning was that this criteria may further enrich our analysis for proteins that directly bind CoA metabolites, since acyl-CoA interaction can underlie both enzymatic and non-enzymatic autoacylation. Using a recently published dataset (Hansen et al., 2019), we identified 6 out of 42 proteins that contain a modified lysine lying in the top 10% of all acetylation stoichiometries measured in HeLa cells (>0.17% stoichiometry, Fig. 5d). Five of these proteins were Uniprot annotated CoA binders (ACLY, CREBBP, HADH, NAT10, NAA10), while one was a member of an AT complex (NAA15). These analyses suggest a multi-pronged approach assessing i) statistically significant multi-ligand competition, ii) dose-response clustering, and iii) acetylation stoichiometry may prove most useful for annotation of the CoA-binding proteome, with the caveat that additional stringency will also lead to filtering of some ‘true’ positives. These findings also imply the acyl-CoA binding proteome interrogatable by this analysis is well-annotated.

Finally, we asked whether CATNIP could provide insight into the function of acyl-CoA binding, particularly whether these events are able to drive non-enzymatic acylation. Our previous studies have found lysine malonylation and succinylation serve as markers of non-enzymatic acylation, due to the high reactivity of their acyl-CoA precursors (Kulkarni et al., 2017). This led us to hypothesize that if an enzyme 1) binds a malonyl-CoA analogue and 2) possesses a malonylation site that overlaps with high stoichiometry acetylation, then acyl-CoA binding may be responsible for driving non-enzymatic acylation. Examining the six proteins above, only one (NAT10) exhibited statistically significant competition by the malonyl-CoA surrogate acetic-CoA. In line with this, while 5/6 of these proteins were found to harbor sites of lysine malonylation (Colak et al., 2015), only in the case of NAT10 were the high stoichiometry acetylation and malonylation sites found on the same residue (K426). This lysine lies within NAT10’s GNAT domain and is highly conserved from eukaryotes to bacteria (Fig. S5f). Analyzing the position of K426 using the structure of a NAT10 orthologue shows it lies proximal to the acetyl-CoA binding site (Chimnaronk et al., 2009), potentially priming it for non-enzymatic acetylation (Fig. 5f). Consistent with this, we find FLAG-NAT10 overexpressed in HEK-293 cells is readily malonylated upon incubation with malonyl-CoA (Fig. 5g; note that we used malonyl- and not acetyl-CoA to decouple enzymatic and non-enzymatic mechanisms, as no known ATs use malonyl-CoA as a cofactor). Such a non-enzymatic mechanism would reconcile the paradoxical finding that this enzyme has been found to undergo functional lysine acetylation in its active site (Cai et al., 2017), but its only biochemically validated catalytic substrates are RNA cytidine residues (Ito et al., 2014). Further work will be needed to evaluate the impact of K426 malonylation on NAT10 activity, as well as the global extent to which malonylation is dependent on specific malonyl-CoA/protein interactions as compared to stochastic labeling of solvent accessible lysine residues. These studies demonstrate the potential for interfacing CATNIP datasets with analyses of lysine malonylation to identify proximity-dependent non-enzymatic acylation mechanisms.

## Discussion

Chemoproteomics has recently emerged as a powerful method for the interrogation of metabolite signaling. Here we describe the development and application of CATNIP, a systems chemoproteomic approach for the high-throughput analysis of acyl-CoA/protein interactions. We first validate the ability of CATNIP to identify protein subsets enriched in CoA-binding, and then apply this method to probe the selectivity of acyl-CoA/protein interactions, visualize novel acyl-CoA interactive biological networks, and characterize the interplay between direct acyl-CoA binding and covalent lysine acylation. CATNIP identified a strong interaction of the RNA cytidine acetyltransferase NAT10 with the feedback metabolite CoA as well as several additional acyl-CoA cofactors. Furthermore, in cell models where acetyl-CoA biosynthesis is impaired we found that a subset of cytidine acetylation in was decreased, implying these CoA metabolites may be capable of interacting with NAT10 as endogenous inhibitors. Of note, the percentage of relative abundance of ac4C is ~8-fold lower in oligo(dT)-enriched RNA than total RNA, and no data regarding the stoichiometry of these targets has been reported. Therefore, additional work will be needed to validate this finding, as well as to understand what effect acetyl-CoA metabolism has on the acetylation of specific RNA targets and pathogenic NAT10 activity. These studies highlight the ability of CATNIP to identify biological processes conditionally regulated by acetyl-CoA and provide a novel hypothesis generation tool for the functional interrogation of metabolite-protein interactions in biology and disease.

To explore the utility of CATNIP for discovery applications, we developed an unbiased workflow to applying chemoproteomic data for the de novo annotation of acetyl-CoA binding proteins. Critical to this endeavor was the integration of CATNIP and acetylation stoichiometry datasets (Hansen et al., 2019), which allowed the identification of a protein subset highly enriched in CoA-binders and AT interactors that was obscure to either method alone (Fig. S4d). An interesting finding was the absence of any ‘unexpected interactors,’ i.e. unannotated proteins with CATNIP profiles indicative of CoA-binding, within this highly curated subset. This suggests the current CoA-binding proteome is well-annotated, with the caveat that this conclusion is entirely dependent on the unique workflow applied here, and therefore does not preclude the discovery of novel acyl-CoA-binding proteins by new experimental methods (e.g. structurally distinct capture probes) or computational analyses. With regards to the latter, it is important to note that many authentic acyl-CoA-binding proteins sampled by CATNIP do not exhibit high stoichiometry acetylation sites (e.g. ATAT1) or fall outside of dose-dependent cluster 1 (e.g. KAT2A). Our studies demonstrate how acetylation stoichiometry may serve as a useful guide to high-confidence annotation of acyl-CoA binding, while simultaneously raising the possibility of mining additional CoA binders and AT interactors from CATNIP data.

Acyl-CoA/protein interactions can play many potential functional roles (Fig. 1). Inspired by recent chemoproteomic studies showing that inositol polyphosphate binding can trigger non-enzymatic protein pyrophosphorylation (Wu et al., 2016), we wondered whether acyl-CoA binding may similarly be a major driver of non-enzymatic lysine acylation. Examining the lysine malonylation, a putative non-enzymatic PTM derived from the electrophilic metabolite malonyl-CoA (Kulkarni et al., 2017), we identified NAT10 as a unique case in which these PTMs could be correlated with proximity to an acyl-CoA binding site. However, this approach is far from predictive and, even in our curated dataset of high confidence acyl-CoA-binding proteins, found many sites of malonylation mapping far from the annotated active site (Fig. 5e, Fig. S5f) (Colak et al., 2015). Although additional studies are needed, our data suggests for many non-enzymatic acylations factors independent of acyl-CoA binding affinity such as lysine nucleophilicity, surface accessibility, and exposure to high local concentrations of electrophilic CoAs may be important determinants for covalent modification.

Finally, it is important to note some limitations of our current method, as well as steps that may be taken to optimize it for future applications. To facilitate the development of CATNIP, our initial study employed ion trap mass spectrometers for protein identification. For future experiments, we propose using higher resolution instruments to simultaneously perform and PTM identification such as lysine acetylation on enriched proteins, which may be indicative of activity, or use tandem-mass tag (TMT) workflows that enable multiplexed measurements in a single LC-MS/MS run. Transitioning CATNIP to higher resolution instruments will be important for improving the throughput and quantitative applications of our method. An important characteristic of CATNIP is that it reports on relative, rather than absolute, binding affinities due to differences in the inherent binding affinity of individual proteins to the Lys-CoA capture matrix. This means CATNIP is best suited to gauging the comparative pharmacology of individual acyl-CoA binding proteins (i.e. for a series of ligands, which ones interact strongly with protein of interest), rather than rank order comparisons of absolute ligand-protein binding affinity across the proteome. Such biases are an intrinsic feature of chemoproteomic methods and extend even to label-free approaches such as LiP-MS and CETSA (Piazza et al., 2018; Sridharan et al., 2019), whose detection of protein-ligand interactions require ligand binding to alter proteolytic or thermal stability, respectively. Future studies of acyl-CoA/protein interactions will likely benefit from the integration of multiple approaches. Spike-in controls whose affinity for the CATNIP matrix has been determined may also prove useful for quantitative measurements. Clustering analysis indicated that many CoA binders and AT interactors display similar competition profiles, implying CATNIP as currently constituted is not able to discriminate between direct and indirect interactors. In addition to using acetylation stoichiometry as an orthogonal measure for the de novo assignment of direct acyl-CoA binding, it may be possible to distinguish indirect binding based on susceptibility to ionic competition (i.e. high salt) or by complementing matrix-based pulldown with covalent capture using clickable photoaffinity probes (Montgomery et al., 2014). Alternatively this may be solved by optimized computational analysis, in which the proteins identified from multiple competitive ligands are compared using topological scoring (TopS) (Sardiu et al., 2019) to determine enrichment of proteins and direct interactions from a range of concentrations or ligand types. Although we focused here on studying the interactions of proteins with endogenous acyl-CoA metabolites, recently multiple classes of drug-like KAT inhibitors have been reported (Baell et al., 2018; Lasko et al., 2017), and we anticipate our method will be immediately useful for understanding the pharmacological specificity and potency of these small molecule chemical probes. Such studies are underway, and will be reported in due course.

## Significance

Acyl-CoAs are essential for life. These central metabolites interact with proteins to regulate critical biological processes ranging from energy production to gene expression. However, despite their importance, the scope and selectivity of these interactions remains unknown. Here we report the development of CATNIP (<u>C</u>oA/<u>A</u>cetyl<u>T</u>ra<u>N</u>sferase <u>I</u>nteraction <u>P</u>rofiling), an approach that combines chemoproteomic profiling with systems-level analysis for the proteome-wide interrogation of acyl-CoA/protein interactions. We validated the ability of this approach to identify CoA-utilizing enzymes based on their dose-dependent metabolite competition profiles and detect unique interaction signatures of enzymes with acyl-CoAs responsible for substrate utilization or metabolic inhibition. Applying CATNIP to profile the susceptibility of acetyltransferases led to the characterization of the metabolic regulation of the RNA acetyltransferase NAT10. Finally, we demonstrated an unbiased workflow for analysis of CATNIP binding data that can be used to detect new candidate acyl-CoA regulated protein networks, identify proteins that directly bind acetyl-CoA, and define proteins whose functional interaction with acyl-CoAs may lead to non-enzymatic acylation. Our studies illustrate the utility of integrating chemical biology and systems biology to rapidly characterize protein-metabolite interactions and provide a powerful first-in-class resource for studying the signaling functions of acyl-CoAs. Moreover, our experimental demonstration that acyl-CoAs show selectivity in their interactions with the AT superfamily suggests manipulating them may re-direct the activity of specific enzyme subsets, and provides a rationale for the development of new approaches to modulate acyl-CoA metabolism in cells and living organisms.

## Author Contributions

Conceptualization, M.J.L., M.P.W., M.E.S., D.C.M., and J.L.M.; Methodology, M.J.L., M.E.S., L.F., M.P.W., and J.L.M..; Data Analysis and Curation, M.J.L., M.E.S., S.D.F., Q.L., and J.L.M.; Investigation and Validation, M.J.L., D.C.M., M.E.S., S.E.B., K.N., J.M., and A.L.T.; Resources, S.D.F., Q.L., and T.A..; Writing – Original Draft, J.L.M.; Writing – Review & Editing, M.J.L., L.F., M.P.W., J.M. and J.L.M.; Supervision and funding acquisition, M.P.W. and J.L.M.

## Supplemental Information

Additional data including Figures S1–S5, Tables S1-S8, and experimental protocols are available in the methods section at the end of this file and accompanying supplemental information.

## Acknowledgements

The authors thank K. Wellen (University of Pennsylvania) and N. Snyder (Drexel University) for helpful discussions. This work was supported by the Intramural Research Program of the NIH, National Cancer Institute, Center for Cancer Research (ZIA BC011488–04), the Stowers Institute for Medical Research, and the National Institute of General Medical Sciences of the National Institutes of Health under Award Number RO1GM112639 to MPW. In addition, this project has been funded in whole or in part with Federal funds from the National Cancer Institute, National Institutes of Health, under contract number HHSN261200800001E. The content is solely the responsibility of the authors and does not necessarily represent the official views of the National Institutes of Health.

## Supplemental Figures

**Figure S1.**
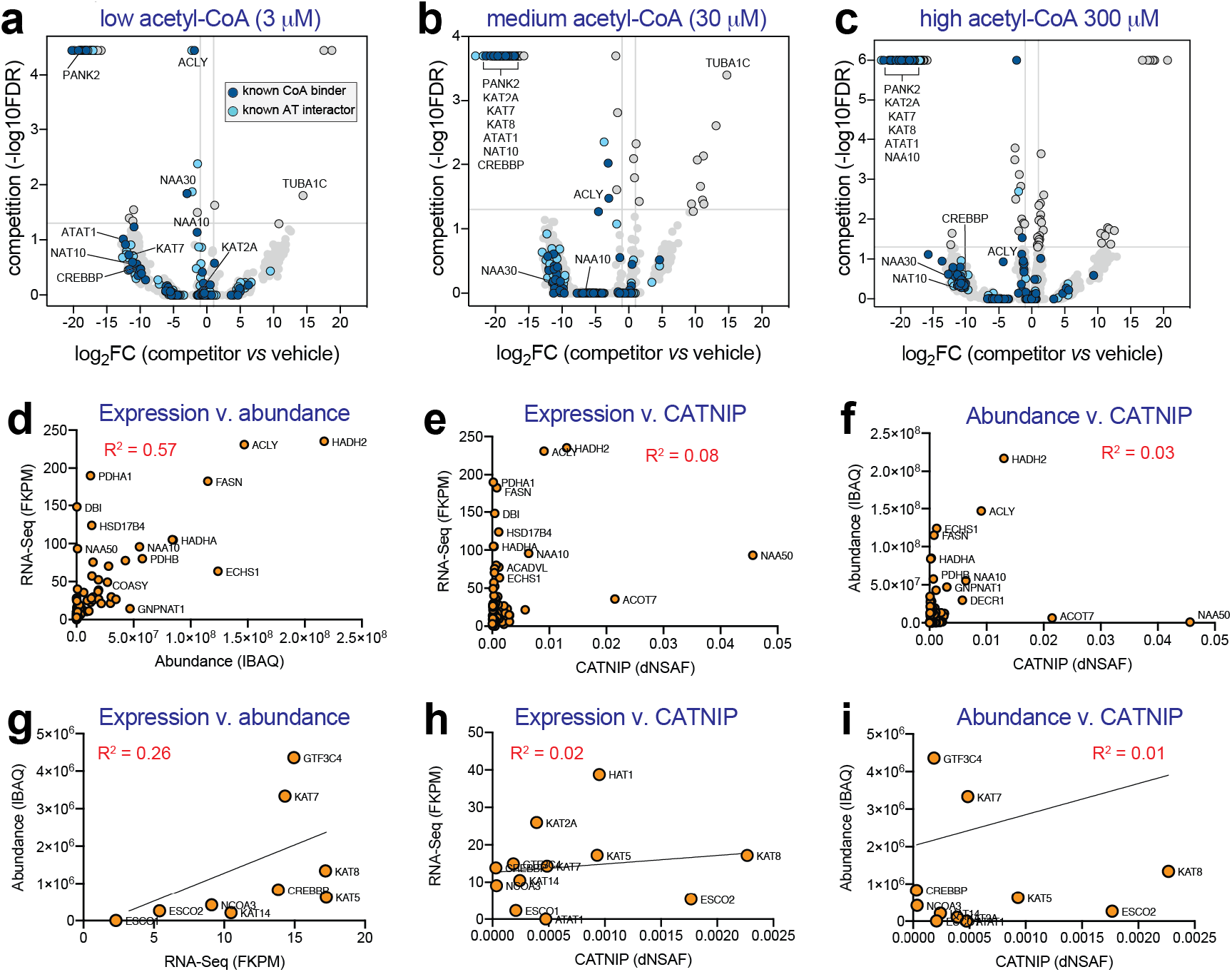
(a-c) Volcano plots depicting competition of proteins captured by Lys-CoA Sepharose by increasing concentrations of acetyl-CoA (0.003, 0.03, and 0.3 μM). Uniprot-annotated CoA-binding proteins (“CoA binder”) as well as members of acetyltransferase complexes (”AT interactors”) are highlighted in dark blue and light blue, respectively. (d-f) Correlation of CATNIP capture efficiency of CoA-binders with gene expression by RNA-Seq (‘expression’) or protein abundance by proteomics (‘abundance’). (g-i). Correlation of CATNIP capture efficiency of annotated ATs with gene expression by RNA-Seq (‘expression’) or protein abundance by proteomics (‘abundance’).

**Figure S2.**
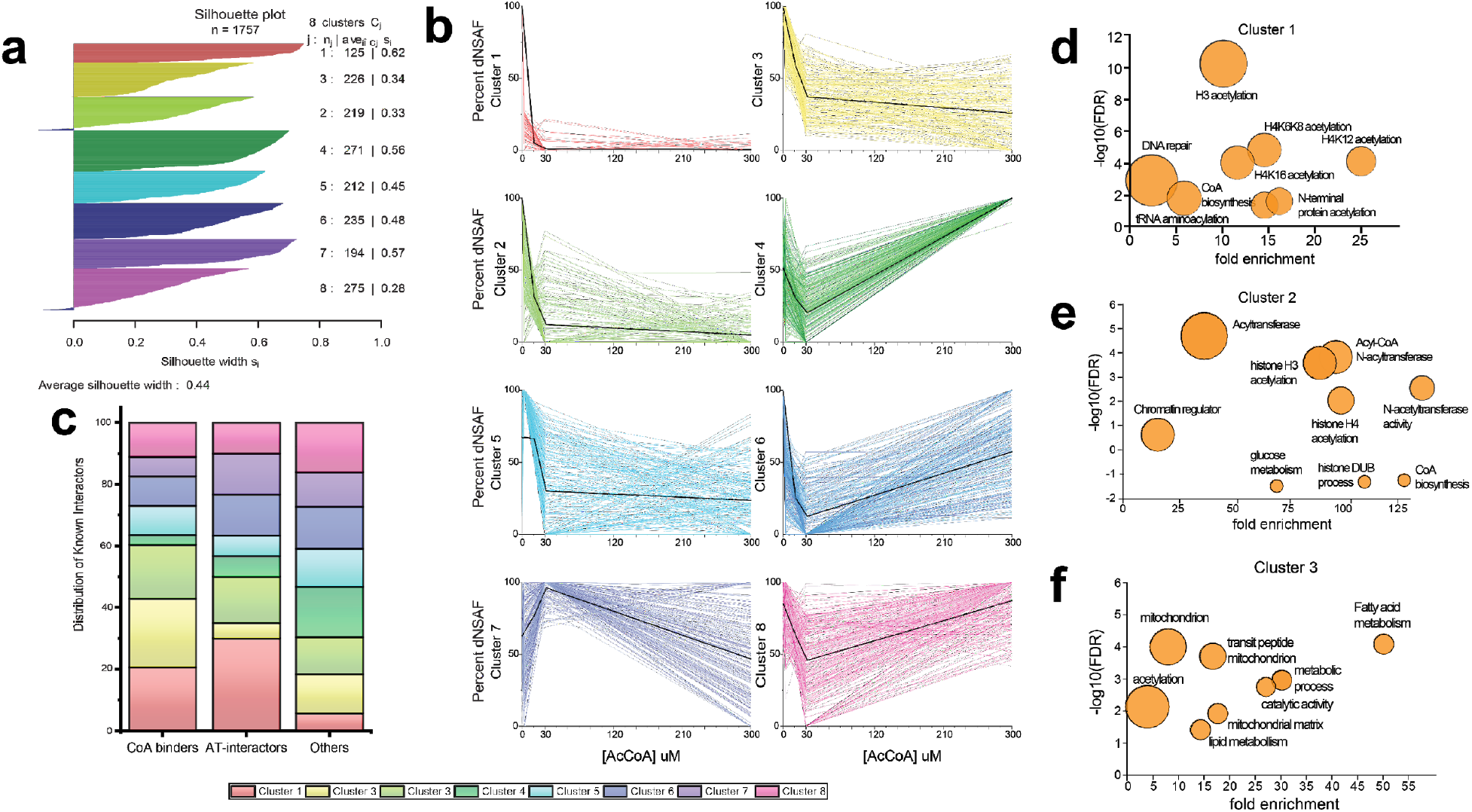
(a) Silhouette plot representative of the 1757 proteins generated in R demonstrating fit of each cluster to the data. (b) Dose-response profiles for all 8 CATNIP clusters (cluster 7 and 8 were not depicted in Fig. 2). Colored lines indicate the capture profiles of individual proteins within each cluster in the presence of increasing concentrations of acetyl-CoA competitor. Black lines indicate the mean capture profile for all proteins in a given cluster. (c) Distribution of CoA-binders, AT-interactors, and non-annotated (‘other’) proteins in each cluster. Clusters 1-3 contain the majority of annotated CoA-binders and AT-interactors, while non-annotated proteins are distributed fairly evenly between different dose-response profiles. (d-f) Gene ontology analysis of Uniprot annotated CoA-binding and AT interacting proteins lying in CATNIP clusters 1-3. Fold enrichment of a specific functional term is plotted versus statistical significance (-log10[FDR]). The circle size reflects the number of proteins matching a given term. Functional enrichment was performed with the tool DAVID (https://david.ncifcrf.gov) by using GO and Swiss-Prot Protein Information Resource terms.

**Figure S3.**
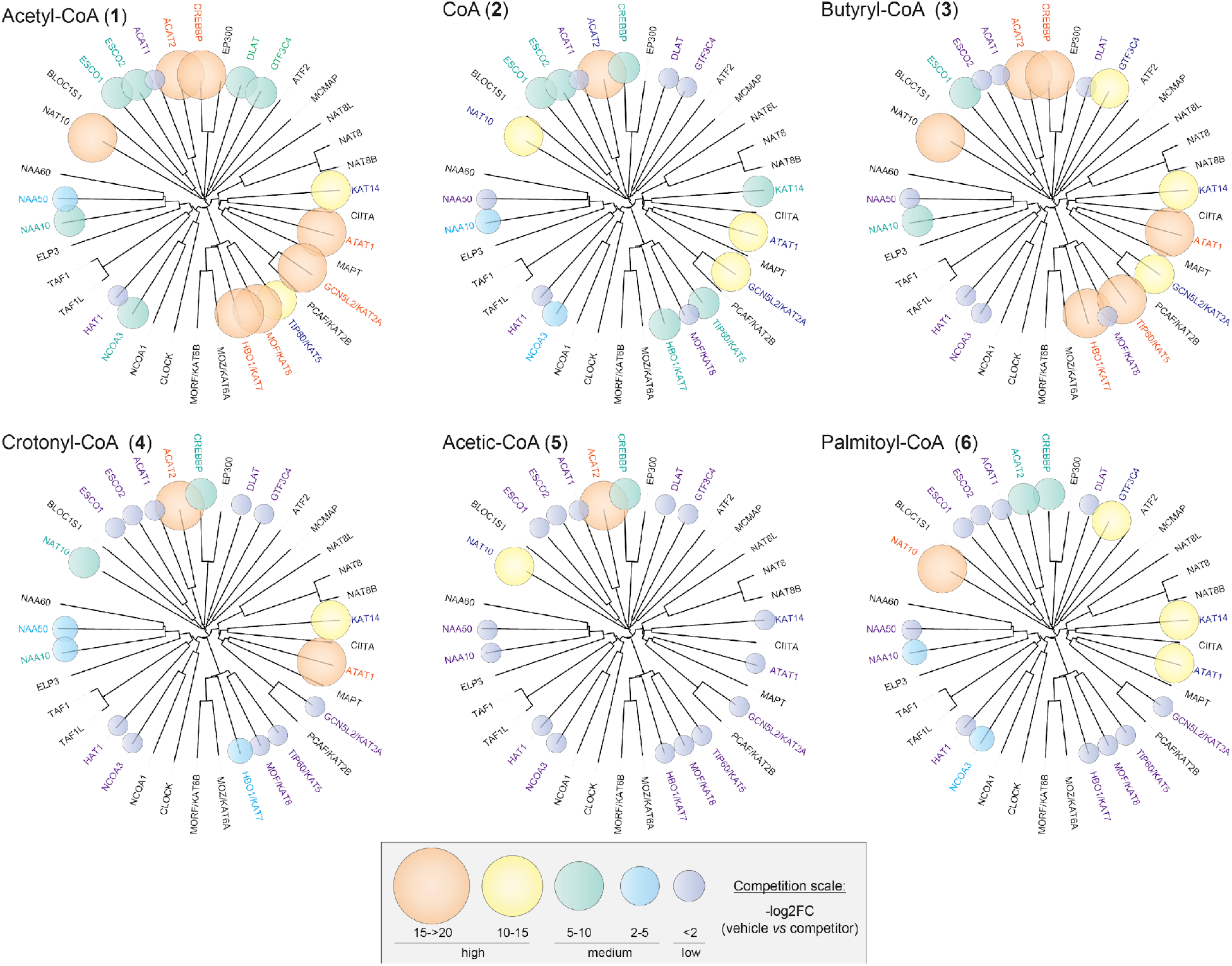
Metabolic acyl-CoA competition profiles of the KAT superfamily. For each protein, metabolic acyl-CoA competition data was overlaid on the unrooted phylogenetic tree of the KAT superfamily. Bubble size and color corresponds to degree of competition observed when proteomes were pre-incubated with CoA metabolites **1**-**6**. All ligands were assessed at 30 μM besides **1**-**2**, which were assessed at both 3 and 30 μM, and **6** whose competition assessed only at 3 μM. In cases where competition was observed at multiple concentrations, the larger log-transformed fold-change (-log2FC) value was used for graphical depiction.

**Figure S4.**
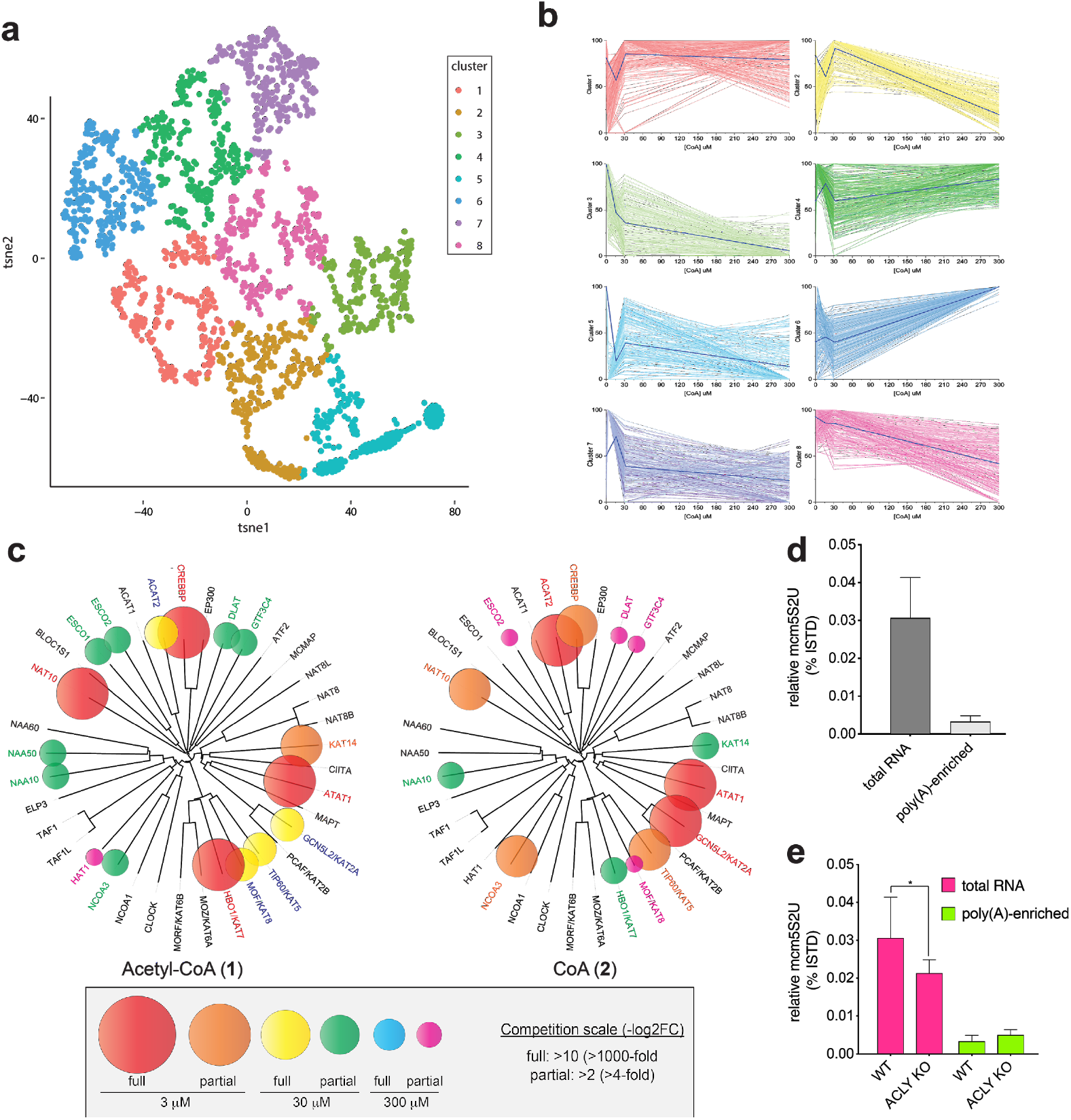
Profiling CoA/protein interactions using CATNIP. (a) t-SNE plot of proteins competed at various concentrations of CoA with proteins colored by the cluster number. Eight clusters were identified as optimal by k-means analysis. Lys-CoA Sepharose was incubated with proteomes in the presence of increasing concentrations of CoA (0, 3, 30, and 300 μM) and protein enrichment was quantified based on distributed normalized spectral abundance factor (dNSAF). Dose-response profiles were transformed into two-dimensional data and used for clustering analysis. n=3 LC-MS/MS runs for each condition. (b) Dose-response profiles of CoA CATNIP clusters. Colored lines indicate the capture profiles of individual proteins within each cluster in the presence of increasing concentrations of CoA competitor. Dark blue lines indicate the mean capture profile for all proteins in a given cluster. (c) Comparing acetyl-CoA and CoA competition of KAT superfamily capture. Circle colors indicate the lowest concentration at which acetyl-CoA (left) or CoA (right) caused a greater than two-fold log-transformed fold change (-log2FC) in the capture of each of each KAT by Lys-CoA Sepharose, while the size of the oval indicates the degree of competition. (d) Monitoring levels of the transfer RNA (tRNA) modification 5-methoxycarbonylmethyl-2-thiouridine (mcm5S2U) indicates efficient depletion of tRNAs during sequential poly(A)-enrichment steps using oligo(dT) beads. (e) Levels of the Elp3 AT-dependent RNA modification mcm5S2U are sensitive to ACLY KO in total RNA but not in poly(A) fraction. Of note, NAT10 AT-dependent ac4C displays the opposite profile (Fig. 4e). Values represent ≥ 3 replicates, analyzed by two-tailed student’s t-test (ns = not significant, *P<0.05, **P<0.01, ***P<0.001).

**Figure S5.**
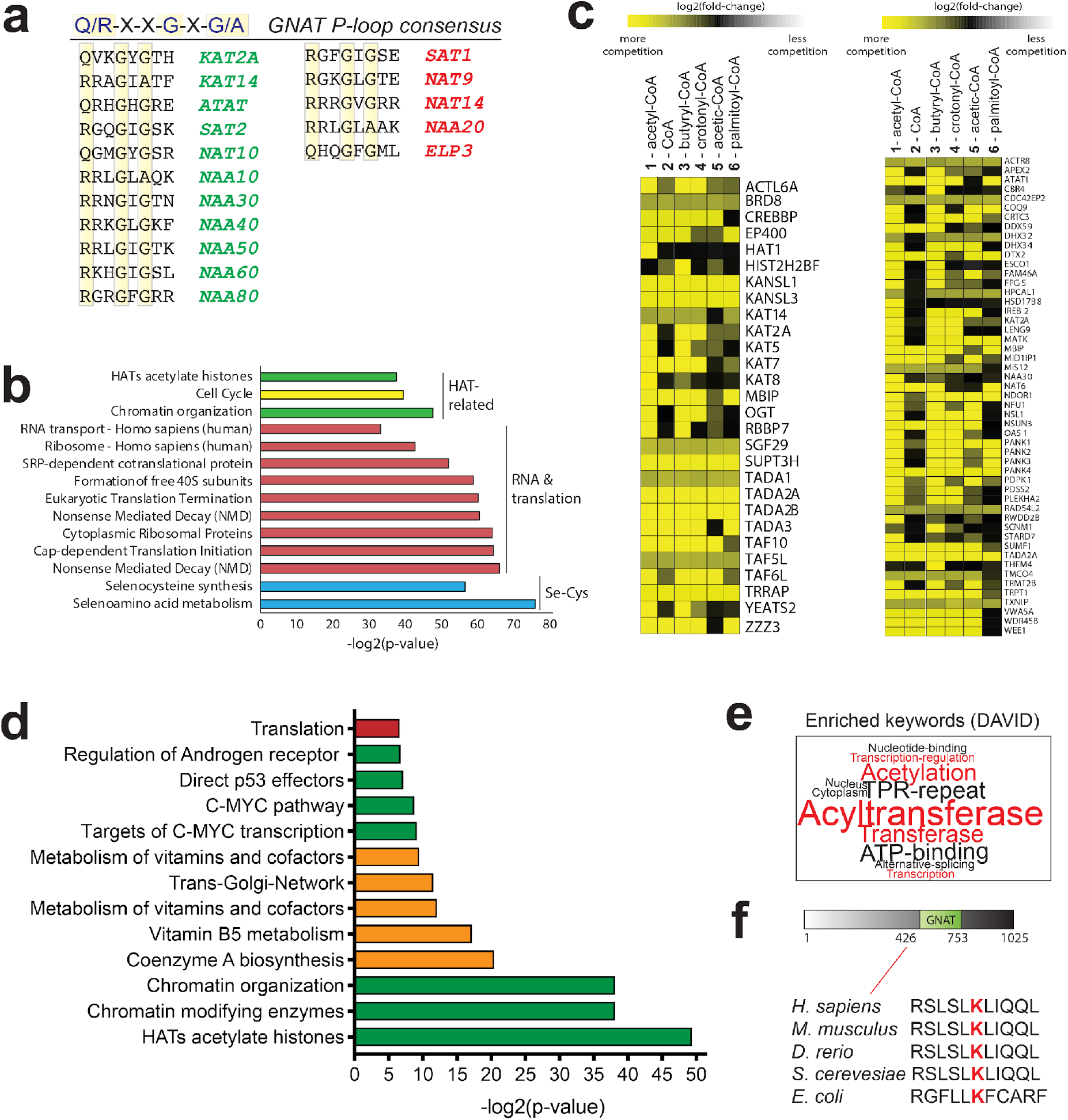
(a) Comparison of GNAT consensus P-loop sequence in proteins captured (left, green) or not captured (right red) by CATNIP. (b) Pathway analysis of proteins identified as potential acyl-CoA interactors by de novo filtering of CATNIP data. Enriched pathways were identified by ConsensusPathDB analysis of 672 proteins displaying multi-ligand (>3) competition and absence (<4 spectral counts) in common contaminant (CRAPome) database. Terms are grouped according to biological functions. (c) Comparing multi-ligand competition profiles of proteins lying within pathways enriched by de novo CATNIP dataset. Left: the 28 proteins enriched in “HATs acetylate histones” pathway. Right: Potentially novel interactors not detected in the CrapOme database. Yellow = more competition of capture, black = less competition of capture. (d) Pathway analysis of proteins identified as potential acyl-CoA interactors by de novo filtering of CATNIP data and combined with dose-dependent acetyl-CoA CATNIP clustering. Enriched pathways were identified by ConsensusPathDB analysis of 45 proteins displaying multi-ligand (>3) competition, absence (<4 spectral counts) in common contaminant (CRAPome) database, and hypersensitivity to dose-dependent acetyl-CoA competition (cluster 1, Fig. 2b-c). (e) Word cloud depicting enriched keywords identified by DAVID analysis of protein subset defined in S5d. Word size corresponds to fold-enrichment of each keyword. (f) NAT10 K426 is conserved from eukaryotes to bacteria.

## Materials and Methods (STAR Methods)

### General materials and methods

NHS-Activated Sepharose 4 Fast Flow resin was purchased from GE Healthcare Life Sciences (71-5000-14 AD). Amine-functionalized Lys-CoA-Ahx was synthesized as described previously (Montgomery et al., 2016). U-^13^C6-glucose (CLM-1396) and U-^13^C2-acetate (CLM-440) were purchased from Cambridge Isotope Laboratories. Acetyl-CoA (A2056), butyryl-CoA (B1508), crotonyl-CoA (28007) and palmitoyl-CoA (P9716) were purchased from Sigma. Coenzyme A (CoA; C7505-51) was purchased from United States Biological. Acetic-CoA was synthesized from 2-bromoacetic acid and CoA in a single-step as described previously (Montgomery et al., 2014). Prior to utilization all acyl-CoAs were analyzed for purity by LC-MS and re-purified via HPLC if necessary. CoAs were quantified using the molar extinction coefficient (ε) for Coenzyme A of 15, 000 M^−1^cm^−1^ at λ_max_ of 259 nm. HeLa cells used to prepare proteomic extracts were grown by Cell Culture Company (formerly National Cell Culture Center, Minneapolis, MN). Proteomes were prepared by re-suspending cell pellets in ice-cold PBS containing protease inhibitor cocktail (1×, EDTA-free, Cell Signaling Technology # 5871S). Samples were then lysed by sonication using a 100 W QSonica XL2000 sonicator (3 × 1 s pulse, amplitude 1, 60 s resting on ice between pulses). Lysates were pelleted by centrifugation (20,817 r.c.f. x 30 minutes, 4 °C) and quantified using the Qubit 4.0 Fluorometer and Qubit Protein Assay Kit. Quantified proteomes were diluted and stored at −80 °C in between uses. TRIzol reagent (#15596026) and oligo-(dT)_25_ Dynabeads (#61005) were purchased from ThermoFisher Scientific (15596026). Analytical analyses of Lys-CoA and all acyl-CoAs were performed using a Shimadzu 2020 LC-MS system.

### Preparation of Lys-CoA Sepharose resin

Lys-CoA Sepharose (**1**) was prepared using NHS-Activated Sepharose 4 Fast Flow resin essentially according to the manufacturer’s protocol (GE Healthcare Life Sciences, Instructions 71-5000-14 AD) (Montgomery et al., 2016). Briefly, amine-functionalized Lys-CoA-Ahx was prepared as a 3.4 mM solution in PBS. Resin was washed with cold 1 mM HCl prior to coupling, before addition of the ligand solution at a ratio of 2:1 resin:ligand volume. The pH was adjusted to ~7-8 by addition of 20x PBS, and the mixture was then rotated at 4°C overnight. The resin was pelleted at 1400 rcf for 3 minutes, and the supernatant was discarded prior to addition of 3 resin volumes of 0.1 M Tris-HCl [pH 8.5], and the mixture was rotated for 3 hr at room temperature. Resin was washed 3x each with alternating solutions of 0.1 M Tris-HCl [pH 8.5] and 0.1M Sodium Acetate, 0.5 M NaCl [pH 4.5] (6 washes total). Resin stored as a 33% solution in aqueous 20% EtOH at 4°C.

### Procedure for CATNIP affinity capture, competition and LC-MS/MS studies

Affinity capture using Lys-CoA Sepharose was carried out essentially as previously reported (Montgomery et al., 2016; Montgomery et al., 2015). Briefly, 33 μl of capture resin was washed once with 1 ml of PBS, prior to addition of 500 μl of clarified lysates (1.5 mg/ml, pretreated with vehicle or competitor ligand for 30 min on ice). This mixture was rotated for 1 hr at room temperature, pelleted at 1400 rcf, and supernatant discarded. Sepharose capture resins were subjected to a series of mild washes using ice cold wash buffer (50 mM Tris-HCl [pH 7.5], 5% glycerol [omitted in LC-MS/MS experiments], 1.5 mM MgCl2, 150 mM NaCl, 3 × 500 μl). Following the last wash, enriched resin was collected on top of centrifugal filters (VWR, 82031-256). For LC-MS/MS analysis of captured proteins, enriched resin was transferred from centrifugal filters to fresh 1.7-ml tubes using 400 μl of tryptic digest buffer (50 mM Tris-HCl [pH 8.0], 1 M urea). Digests were initiated by addition of 0.4 μl of 1 M CaCl2 and 4 μl of trypsin (0.25 mg/ml) and allowed to proceed overnight at 37°C with shaking. After extraction, tryptic peptide samples were acidified to a final concentration of 5% formic acid, lyophilized, and frozen at −80°C until LC-MS/MS analysis.

### MudPIT LC-MS/MS analysis of and database searching of Lys-CoA enriched proteomes

Lyophilized peptide samples from Lys-CoA Sepharose enriched HeLa proteomes were analyzed independently in triplicate by Multidimensional Protein Identification Technology (MudPIT), as described previously (Florens and Washburn, 2006; Washburn et al., 2001a). Briefly, dried peptides were resuspended in 100μL of Buffer A (5% acetonitrile (ACN), 0.1% formic acid (FA)) prior to pressure-loading onto 100 μm fused silica microcapillary columns packed first with 9 cm of reverse phase (RP) material (Aqua; Phenomenex), followed by 3 cm of 5-μm Strong Cation Exchange material (Luna; Phenomenex), followed by 1 cm of 5-μm C_18_ RP. The loaded microcapillary columns were placed in-line with a 1260 Quartenary HPLC (Agilent). The application of a 2.5 kV distal voltage electrosprayed the eluting peptides directly into LTQ linear ion trap mass spectrometers (Thermo Scientific) equipped with a custom-made nano-LC electrospray ionization source. Full MS spectra were recorded on the eluting peptides over a 400 to 1600 *m*/*z* range followed by fragmentation in the ion trap (at 35% collision energy) on the first to fifth most intense ions selected from the full MS spectrum. Dynamic exclusion was enabled for 120 sec (Zhang et al., 2009). Mass spectrometer scan functions and HPLC solvent gradients were controlled by the XCalibur data system (Thermo Scientific).

RAW files were extracted into .ms2 file format (McDonald et al., 2004) using RawDistiller v. 1.0, in-house developed software (Zhang et al., 2011). RawDistiller D(g, 6) settings were used to abstract MS1 scan profiles by Gaussian fitting and to implement dynamic offline lock mass using six background polydimethylcyclosiloxane ions as internal calibrants (Zhang et al., 2011). MS/MS spectra were first searched using ProLuCID (Xu et al., 2015) with a mass tolerance of 500 ppm for peptide and fragment ions. Trypsin specificity was imposed on both ends of candidate peptides during the search against a protein database combining 36,628 human proteins (NCBI 2016-06-10 release), as well as 193 usual contaminants such as human keratins, IgGs and proteolytic enzymes. To estimate false discovery rates (FDR), each protein sequence was randomized (keeping the same amino acid composition and length) and the resulting “shuffled” sequences were added to the database, for a total search space of 73,642 amino acid sequences. A mass of 15.9949 Da was differentially added to methionine residues.

DTASelect v.1.9 (Tabb et al., 2002) was used to select and sort peptide/spectrum matches (PSMs) passing the following criteria set: PSMs were only retained if they had a DeltCn of at least 0.08; minimum XCorr values of 1.9 for singly-, 2.7 for doubly-, and 2.9 for triply-charged spectra; peptides had to be at least 7 amino acids long. Results from each sample were merged and compared using CONTRAST (Tabb et al., 2002). Combining all replicate injections, proteins had to be detected by at least 2 peptides and/or 2 spectral counts. Proteins that were subsets of others were removed using the parsimony option in DTASelect on the proteins detected after merging all runs. Proteins that were identified by the same set of peptides (including at least one peptide unique to such protein group to distinguish between isoforms) were grouped together, and one accession number was arbitrarily considered as representative of each protein group.

*NSAF7* (Zhang et al., 2010) was used to create the final reports on all detected peptides and non-redundant proteins identified across the different runs. Spectral and protein level FDRs were, on average, 0.31±0.10% and 1.0±0.35%, respectively. QPROT (Choi, et al, 2015) was used to calculate a log fold change and false discovery rate for the dosed samples compared to the vehicle control.

### Partitioning clustering

To group proteins based on their abundance profile across the four treatment conditions (i.e. 0, 3μM, 30μM and 300μM), first each individual protein was normalized in each condition to the highest value across the four conditions (i.e. the highest value equals to 100%). To spatially map the proteins in the dataset, a t-distributed stochastic neighbor embedding (t-SNE), a nonlinear visualization of the data was applied. Then, k-means clustering was applied to this transformed matrix using the Hartigan-Wong algorithm and a maximum number of iterations set at 50000. To determine the best partition, the numbers of clusters, k, were continuously increased from 3 to 20. The result showed that the optimal number of clusters was obtained when k=8, after carefully inspecting all the clusters and their silhouette and Hartigan indexes. All computations were run using R environment using k-means function for the partition and daisy function to compute all the pairwise dissimilarities (Euclidean distances) between observations in the dataset for the silhouette.

### Dose response curves

Normalized dNSAF values for each protein were plotted as a function of ligand concentration in Origin Pro 2018 for each cluster. The curves were averaged in Origin and the average was displayed on the graph.

### Topological data analysis

The input data for TDA were represented in a matrix, with each column corresponding to a CoA ligand and each row corresponding to a protein. Values were distributed spectral counts values for each protein. A network of nodes with edges between them was then created using the TDA approach based on Ayasdi platform (AYASDI Inc., Menlo Park CA as described previously (Sardiu et al., 2015). Two types of parameters are needed to generate a topological analysis: First is a measurement of similarity, called metric, which measures the distance between two points in space (i.e. between rows in the data). Second are lenses, which are real valued functions on the data points. Here, Variance Norm Euclidean was used as a distance metric with 2 filter functions: Neighborhood lens 1 and Neighborhood lens2. Resolution 30 and gain 3 were used to generate Fig. 5c.

### Pathway analysis

Proteins that were changing in at least one of the CoA ligands with a Z-score less than −2 and FDR less or equal to 0.05 were considered for the analysis. Using this criteria, 671 proteins were identified and used for the pathway analysis. As expected, HATs acetylate histone was one of the top 30 enriched pathways (p-value of 4.55e-12) in the ConsensusPathDB (http://cpdb.molgen.mpg.de/) database.

### Bioinformatic analyses of CATNIP data and correlation with literature datasets

A list of annotated CoA-binders was defined by searching the Uniprot database using query terms related to this function including “CoA binding,” “CoA,” “Coenzyme A,” “Acetyltransferase” “HAT,” “NAT,” “NAA,” “GNAT.” A similar analysis was performed to annotate AT interactors, using query terms including “HAT complex,” “KAT complex,” “NAA complex,” “NAT complex,” and “acetyltransferase complex.” Results were then manually curated with irrelevant proteins and duplicates removed, resulting in the term list provided in Table S2. Correlation of CATNIP enrichment to HeLa cell gene expression and protein abundance (Fig. S1 d-i) was performed using literature RNA-Seq and deep proteomic datasets (Nagaraj et al., 2011). Venn diagrams comparing overlap between proteins competed 2-fold by acetyl-CoA and all other ligands (Fig. 3b), or metabolic acyl-CoAs **3**-**6** (Fig. 3c) were generated by identifying a list of proteins showing a (-log2FC) value >1 for each ligand and then assessing overlap using an online Venn diagram tool accessible at http://bioinformatics.psb.ugent.be/webtools/Venn/. Protein subsets were interrogated for enrichment of molecular functions and pathways using the online informatics tools DAVID (david.ncifcrf.gov) and ConsensusPathDB (http://cpdb.molgen.mpg.de/CPDB/rlFrame). For analysis of acetylation stoichiometry, filtered protein subsets were cross-referenced with a list of peptide hits falling in the top 10% of all HeLa cell lysine acetylation stoichiometries measured in a recently published analysis (Hansen et al., 2019). For analysis of lysine malonylation, filtered protein subsets were cross-referenced with a list of malonylated peptides derived from a recently published analysis. Figures of *E. coli* NAT10 orthologue complexed with acetyl-CoA was generated using Chimera.

### Isotopic tracing experiments to determine metabolic source of N4-acetylcytidine (ac4C)

HeLa cells were cultured at 37 °C under 5% CO_2_ atmosphere in a growth medium of DMEM supplemented with 10% FBS and 2 mM glutamine. HeLa cells were plated in 10 cm dishes (3 x10^6^ cells in 10 ml RPMI media/dish) and allowed to adhere for 24 h. After this, media was removed, cells were washed once with PBS (10 ml), and switched to either i) heavy glucose media (glucose-free DMEM containing 2 mM glutamine, 25 mM U-^13^C_6_-glucose, 0.2 mM acetate), ii) heavy acetate media (glucose-free DMEM containing 2 mM glutamine, 25 mM glucose, 0.2 mM U-^13^C_2_-acetate) or iii) regular glucose media (glucose-free DMEM containing 2 mM glutamine, 25 mM glucose, 0.2 mM acetate). Cells were incubated with the tracer for 16 h or 24 h at 37 °C and total RNA was harvested using TRIzol reagent (ThermoFisher Scientific) according to the manufacturer’s instructions. Digestion of total RNA (220 μg) was performed as previously described (Sinclair et al., 2017). Briefly, RNA was incubated with 1U/10 μg RNA of nuclease P1 (Sigma-Aldrich) in 100 mM ammonium acetate [pH 5.5] for 16 hr at 37 °C. Five microliter of 1 M ammonium bicarbonate [pH 8.3] and 0.5U/10 μg RNA of Bacterial Alkaline Phosphatase (ThermoFisher Scientific) were added for 2 hr at 37 °C. Following digestion, sample volumes were adjusted to 150 μL with RNase-free water and spin filtered to remove enzymatic constituents (Amicon Ultra 3K, #UFC500396). Filtrate and washes (200 μL x 3, RNase-free water) were collected and lyophilized. Lyophilized samples were reconstituted in 250 μL H2O containing internal standards (D3-ac4C, 500 nM; 15N3-C, 5 ìM, Cambridge Isotopes). Individual samples (15 μL for ac4C analyses, 5 μL for major bases) were then analyzed via injection onto a C_18_ reverse phase column coupled to an Agilent 6410 QQQ triple-quadrupole LC mass spectrometer in positive electrospray ionization mode (Agilent Technologies). Quantification was performed based on nucleoside-to-base ion transitions using standard curves of pure nucleosides and stable isotope labelled internal standards.

### LN229 ac4C analysis and MS

LN229 wild-type (WT) and ACLY knockout (ACLY KO) cell lines (kind gift of K. Wellen laboratory, University of Pennsylvania) were cultured at 37 °C under 5% CO_2_ atmosphere in a growth medium of RPMI supplemented with 10% FBS and 2 mM glutamine as previously described (Lee et al., 2018). For assessment of ac4C levels, total RNA was isolated from LN229 cells using TRIzol reagent (ThermoFisher Scientific). Enrichment of polyadenylated RNA [poly(A) RNA] for UHPLC-MS, was carried using two rounds of selection with Oligo-(dT)_25_ Dynabeads (ThermoFisher Scientific) according to the manufacturer’s instructions. 300 ng of total or poly(A) RNA was used to evaluate the levels of ac4C and mcm5s2U by LC-MS/MS using a similar method as described.^57^ Briefly, prior to UHPLC-MS analysis, 300 ng of each oligonucleotide was treated with 0.5 pg/μl of internal standard (IS), isotopically labeled guanosine, [^13^C][^15^N]-G. The enzymatic digestion was carried out using Nucleoside Digestion Mix (New England BioLabs) according to the manufacturer’s instructions. Finally, the digested samples were lyophilized and reconstituted in 100 μl of RNAse-free water, 0.01% formic acid prior to UHPLC-MS/MS analysis. The UHPLC-MS analysis was accomplished on a Waters XEVO TQ-S™ (Waters Corporation, USA) triple quadruple mass spectrometer equipped with an electrospray source (ESI) source maintained at 150 °C and a capillary voltage of 1 kV. Nitrogen was used as the nebulizer gas which was maintained at 7 bars pressure, flow rate of 500 l/h and at temperature of 500°C. UHPLC-MS/MS analysis was performed in ESI positive-ion mode using multiple-reaction monitoring (MRM) from ion transitions previously determined for ac4C and mcm5s2U (Basanta-Sanchez et al., 2016). A Waters ACQUITY UPLC™ HSS T3 guard column, 2.1x 5 mm, 1.8 μm, attached to a HSS T3 column, 2,1×50 nm, 1.7 μm were used for the separation. Mobile phases included RNAse-free water (18 MΩcm^−1^) containing 0.01% formic acid (Buffer A) and 50:50 acetonitrile in Buffer A (Buffer B). The digested nucleotides were eluted at a flow rate of 0.5 ml/min with a gradient as follows: 0-2 min, 0-10%B; 2-3 min, 10-15% B; 3-4 min, 15-100% B; 4-4.5 min, 100 %B. The total run time was 7 min. The column oven temperature was kept at 35oC and sample injection volume was 10 ul. Three injections were performed for each sample. Data acquisition and analysis were performed with MassLynx V4.1 and TargetLynx. Calibration curves were plotted using linear regression with a weight factor of 1/x.

### Overexpression, non-enzymatic malonylation, and immunoprecipitation of NAT10

HEK-293T cells were plated in 10 cm dishes (2.5 x106 cells/dish in 10 ml DMEM media) and allowed to adhere and grow for 24 h. 3xFLAG-tagged NAT10 was overexpressed using FuGENE^®^ 6 transfection reagent (Promega #E2691) according to the manufacturer’s instructions. Overexpression was carried out by incubating the cells for 24 h at 37°C under 5% CO2 atmosphere, after which time the cells were harvested, and lysed in potassium phosphate buffer, pH 8, sonicated using a 100 W QSonica XL2000 sonicator (3 × 1 s pulse, amplitude 1, 60 s resting on ice between pulses), and quantified using the Qubit Broad Sensitivity Protein Kit (Thermo Fisher # Q33211). Lysates were incubated with 0 or 0.25 mM malonyl-CoA for six hours at 37°C. Anti-FLAG pulldown was performed using FLAG immunoprecipitation kit (Sigma - FLAGIPT1-1KT) according to the manufacturer’s instruction. 450ug of lysate was incubated with the anti-FLAG resin over night at 4°C. Eluted protein was ran on SDS-PAGE and immunoblotted against anti-FLAGtag and anti-Malonyl-Lysine antibodies. For immunoblotting, SDS–PAGE gels were transferred to nitrocellulose membranes (Novex, Life Technologies # LC2001) by electroblotting at 30 V for 1 h using a XCell II Blot Module (Novex). Membranes were blocked using StartingBlock (PBS) Blocking Buffer (Thermo Scientific) for 30 min and incubated overnight at 4 °C in primary antibody. The membranes were washed with TBST buffer and incubated with secondary HRP-conjugated antibody (Cell Signaling #7074) for 1 h at room temperature. The membranes were again washed with TBST and treated with chemiluminescence reagents (Western Blot Detection System, Cell Signaling) for 1 min, and imaged for chemiluminescent signal using an ImageQuant Las4010 Digital Imaging System (GE Healthcare).

### Data accessibility

The mass spectrometry proteomics data have been deposited to the ProteomeXchange Consortium via Pride (Deutsch, et. al, 2017; Perez-Riverol, Y., et. al, 2019) partner repository with the dataset identifier PXD013157 and 10.6019/DXD013157. Original data underlying this manuscript may also be accessed after publication from the Stowers Original Data Repository at http://www.stowers.org/research/publications/libpb-1355. Review access can be obtained using the following username and password:

*Username*

*Password*: 3npaE9w9

